# SARS-CoV-2 leverages airway epithelial protective mechanism for viral infection

**DOI:** 10.1101/2022.01.29.478335

**Authors:** Allison M. Greaney, Micha S.B. Raredon, Maria P. Kochugaeva, Laura E. Niklason, Andre Levchenko

**Affiliations:** Department of Biomedical Engineering, Yale University, New Haven, CT 06511, USA; Vascular Biology and Therapeutics Program, Yale School of Medicine, New Haven, CT 06511, USA; Medical Scientist Training Program, Yale University, New Haven, CT 06511, USA; Yale Systems Biology Institute, Yale University, West Haven, CT 06516, USA; Department of Anesthesiology, Yale School of Medicine, New Haven, CT 06510, USA; Humacyte Inc., Durham North Carolina 27713, USA

**Keywords:** SARS-CoV-2, COVID-19, viral entry, cell system, viral infection, pulmonary epithelium, epithelial defense

## Abstract

Despite much concerted effort to better understand SARS-CoV-2 viral infection, relatively little is known about the dynamics of early viral entry and infection in the airway. Here we analyzed a single-cell RNA sequencing dataset of early SARS-CoV-2 infection in a humanized *in vitro* model, to elucidate key mechanisms by which the virus triggers a cell-systems-level response in the bronchial epithelium. We find that SARS-CoV-2 virus preferentially enters the tissue via ciliated cell precursors, giving rise to a population of infected mature ciliated cells, which signal to basal cells, inducing further rapid differentiation. This feed-forward loop of infection is mitigated by further cell-cell communication, before interferon signaling begins at three days post-infection. These findings suggest hijacking by the virus of potentially beneficial tissue repair mechanisms, possibly exacerbating the outcome. This work both elucidates the interplay between barrier tissues and viral infections, and may suggest alternative therapeutic approaches targeting non-immune response mechanisms.

## Introduction

Since its first outbreak was reported on December 31, 2019 in Wuhan, China(Fan et al., 2020), COVID-19 has claimed over 873,957 lives in the United States, and over 5.58 million globally, so far (Organization, 2022; Prevention, 2022). Despite recent roll-out of vaccines against the virus and a declining case fatality rate in the United States, the pandemic has not abated globally and new, more infectious strains of the virus are still being identified(Chia et al., 2021; Meng et al., 2021; Pulliam et al., 2021). A major challenge to fighting the virus remains the lack of precise characterization of the onset of infection and early disease progression. This limited understanding of the early phases of disease has contributed to the massive loss of life.

The virus, SARS-CoV-2, enters the human host via the ACE2 receptor on ciliated cells of large airways, before spreading locally and systemically (Hoffmann et al., 2020; Hou et al., 2020; Leung et al., 2020; Ravindra et al., 2021; Schaefer et al., 2020; Sungnak et al., 2020). Beyond this, little is known about downstream mechanisms leading to physiologic symptoms, including cough, pneumonia, progression onto ARDS, and death (Arentz et al., 2020; Zhou et al., 2020a).

The initial interactions between airway epithelium and the virus are difficult to evaluate *in vivo*, as patient testing usually occurs in the days following infection. Additionally, the SARS-CoV2 virus has a strong tropism for human cells and is not easily studied in typical animal models. To address these problems, *in vitro* models have been leveraged to observe early stages of infection dynamics in cultured human cells. One common model has utilized cultured human bronchial epithelial cells (HBECs) at an air-liquid interface (ALI). Bronchial cells differentiate into a native-like pseudostratified mucociliary epithelium, which can then be inoculated with virus. These models have proven to be remarkably accurate representations of proximal airway epithelial reaction to SARS-CoV-2 infection (Ravindra et al., 2021; Zhu et al., 2020). They also have the added benefit of avoiding challenges associated with analyzing clinical samples, such as sample degradation and patient-to-patient variability (Huang et al., 2020b; Ravindra et al., 2021).

While the dramatic and unique immunological response to SARS-CoV-2 infection has been well-documented (Giamarellos-Bourboulis et al., 2020; Huang et al., 2020a; Lucas et al., 2020; Mathew et al., 2020; Takahashi et al., 2020; Zhou et al., 2020b), the timing of its initiation is still poorly characterized (Maggi et al., 2020). Indeed, the inflammatory and immune response to the virus may not fully develop in the first few days following infection (Lieberman et al., 2020). These findings raise the question of how the first line of defense, the airway epithelium, responds to the infection over the first few days following exposure, in the absence of a full-fledged immune reaction. Combining biomimetic models of the bronchial epithelium with single-cell RNA sequencing (scRNAseq) analysis methods can provide unprecedented information regarding the dynamic processes of viral infection and host cellular response. Identification and characterization of the cell types that comprise airway epithelium by scRNAseq can lead to better molecular-level understanding of viral tropism, tissue damage, and the dynamics of compensatory tissue reorganization. Furthermore, scRNAseq can provide a detailed view of changing cell-cell communication, reflected in expression patterns of cognate ligand-receptor pairs that are indicative of rapidly evolving cell-cell communication patterns.

In this work, we describe the first three days of SARS-CoV-2 infection in an ALI model of proximal airway epithelium, using a recently published single-cell gene expression dataset(Ravindra et al., 2021). By progressively refining cellular identities, we find that the virus preferentially enters via cells that are in the process of differentiating from basal-secretory intermediate cells or secretory cells into ciliated cells. As these infected ciliated progenitor cells further differentiate following viral infection, the viral infection spreads to more mature ciliated cells over time. Concurrently, viral infection and injury of ciliated epithelium also triggers a rapid enhancement of differentiation of basal cells, perhaps in order to restore the injured ciliated epithelial layer, leading to a substantial transient depletion of the resident progenitor cell populations. By the third day of infection, at the viral load examined, the renewal rate of basal cells is partially restored, thereby replenishing their numbers. These cell population dynamics are correlated with a transient increase of pro-differentiation cues followed by expression of pro-renewal cues for basal cells, which are all secreted by the population of infected ciliated cells. We propose a system-level mechanism that may underlie these processes, and show that this mechanism creates a key vulnerability in airway epithelial tissue innate immunity during the first critical days of viral infection and replication. This work provides a high-resolution and nuanced perspective on dynamics of the proximal epithelial cell system from the onset of SARS-CoV-2 infection, thereby revealing fundamental properties of barrier-tissue response to viral entry and identifying routes for potential targeted interventions.

## Results

### Single-cell analysis of cell population dynamics during SARS-CoV-2 infection

scRNAseq data were obtained from a previous manuscript, wherein ALI cultures of differentiated HBECs (Lonza) were inoculated with SARS-CoV-2 virus and harvested at 1 day post-infection (dpi), 2dpi, and 3dpi(Ravindra et al., 2021). The three infected time points and an uninfected mock condition were sequenced at the single-cell level (Fig. 1A). In a de novo analysis, the scRNAseq data were integrated across the time points (Seurat (Satija et al., 2015)), and all typical proximal airway epithelial cell types were identified, including basal cells, cycling basal cells, a small population of basal cells undergoing EMT (a known artifact of the ALI culture method (Greaney et al., 2020)), intermediate cells (between basal and club phenotypes), club cells, goblet cells, ciliated cells, ionocytes, tuft cells, and pulmonary neuroendocrine cells (PNECs) (Fig. 1B,C). These cell types were identified by top differentially expressed genes (Fig. 1D).

**Figure 1.**
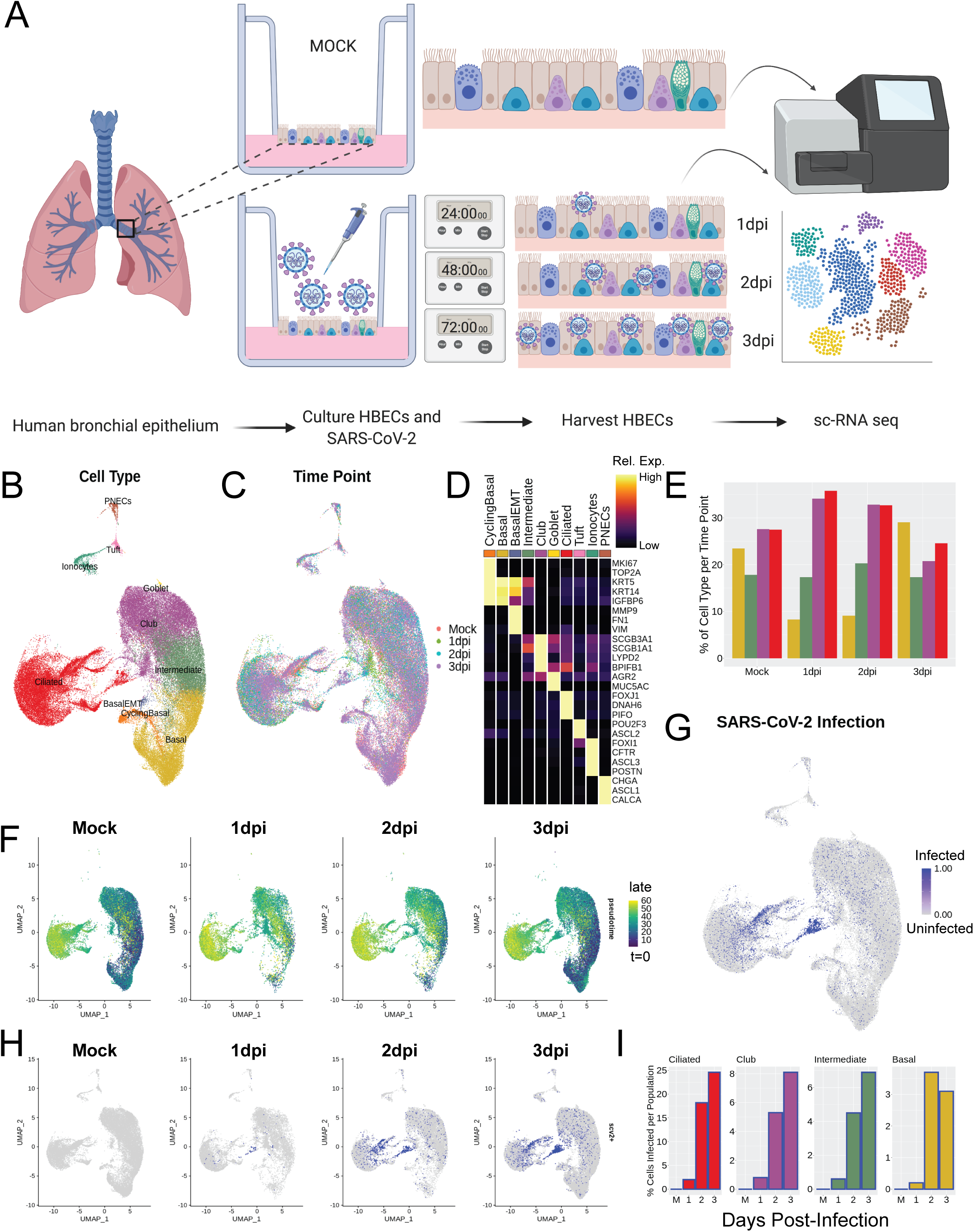
Single-cell analysis of cell population dynamics during SARS-CoV-2 infection. A) Schematic of the experimental protocol; cells under four ALI conditions were collected and transcriptomically sequenced, including the first three days post-viral infection, and an uninfected control condition, denoted as “Mock”. Images adapted from BioRender. B,C) UMAP analysis of scRNAseq data aligned in Seurat, colored and labeled by: (B) cell type and (C) infection condition, to demonstrate batch correction across samples. D) Heatmap of marker genes used for each cell type; columns represent average expression per cluster, rows are unity normalized and colored by relative expression (see Methods). E) Bar graph of cell type proportions per time point for the four largest clusters by cell number: basal, intermediate, club, and ciliated cells. F) UMAP results colored by pseudotime, as calculated using Monocle 3, split by time point, to demonstrate transient loss of progenitor cell populations in early infection timepoints. G,H) UMAP analysis of SARS-CoV-2 infected cells (blue) (G) aligned and (H) split by time point, to track the dynamics of the localized pockets of infected ciliated cells. I) Bar graphs of the proportion of infected cells per cell type at each time point for ciliated, club, intermediate, and basal cells. Cells were identified as infected if they expressed ≥ 10 exonic SARS-CoV-2 viral transcripts (see Methods).

This initial cell type specification was used to study the epithelial cell population dynamics following SARS-CoV-2 infection. Strikingly, we observed a distinct, acute depletion of basal cells at 1dpi, which recovered to near-control (Mock) levels by 3dpi (Fig. 1E). More broadly, at 1dpi, there was a profound depletion of all less-mature cells in the model, as identified by pseudotime (Cao et al., 2019), across all cell types at 1dpi (Fig. 1F). This indicated a transient depletion of all population-specific resident progenitors, suggesting a phenotypic switch from renewal to differentiation, and an increased phenotypic flux toward mature cell types in the infected conditions. Like the basal cell cluster, population-specific resident progenitor cells also recovered their numbers by 3dpi. Using viral transcripts to indicate SARS-CoV-2 infection (consistent with prior methods(Ravindra et al., 2021)), we ascertained that viral load initiates and concentrates in the ciliated cell cluster, but, interestingly, primarily in a subset of cells in one region of the ciliated cluster that express markers of immaturity (Fig. 1G,H). Longitudinal analysis of data on subsequent post-infection days suggests that the viral infection then spreads from this somewhat immature cell type to more mature cells, either by maturation of these infected cells or by viral replication and re-infection of more mature cells. By 3dpi, nearly 25% of ciliated cells in the culture system are infected (Fig. 1I). Overall, our initial observations using dynamic scRNAseq demonstrate a transient depletion of basal and other progenitor cells upon infection, as well as greater SARS-CoV-2 tropism toward an immature subset of ciliated cells.

### SARS-CoV-2 enters via ciliated progenitor cells

To gain a better perspective on SARS-CoV-2 infection dynamics in ciliated cells, we re-analyzed the ciliated cluster to gain a more discrete appreciation of the relevant subtypes (Fig. 2A,B). While all cells in this cluster expressed canonical ciliated cell markers, four ciliated subtypes were identified and named, based on differential gene expression patterns. These subtypes include Ciliated Progenitors, defined by increased basal and secretory markers such as KRT5 and SCGB1A1, respectively; two Mature Ciliated clusters, with Mature Ciliated 1 defined by genes associated with immune response, such as HLA-DRB5, and Mature Ciliated 2 with uniquely high expression of genes associated with cilia motility, such as DLEC1. Finally, we defined a separate subtype of Novel Infected Ciliated cells, which appear only in infected conditions, are not associated with any other clusters, and which express genes unique to viral responsiveness, such as IFIT2 (Fig. 2C).

**Figure 2.**
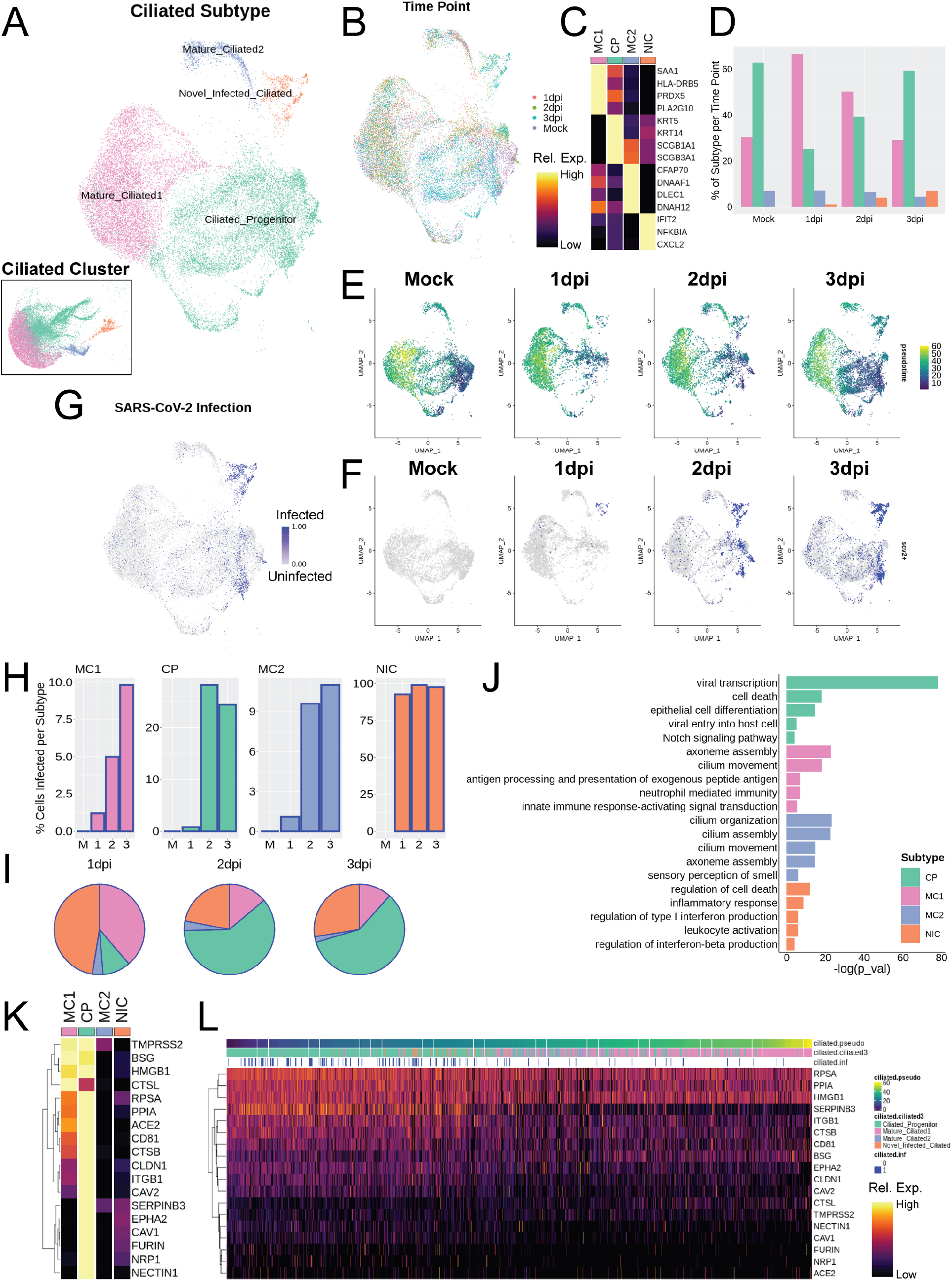
SARS-CoV-2 enters via ciliated progenitor cells. A,B) UMAP of subset ciliated cells colored and labeled by (A) subtype (inset shows subtypes on Fig. 1 embedding) and (B) time point, to demonstrate batch correction across samples. C) Heatmap of marker genes for each ciliated subtype; columns represent average expression per cluster, rows are unity normalized and colored by relative expression (see Methods). D) Bar graph of ciliated subtype proportions per time point. E) UMAP analysis results colored by pseudotime, as calculated using Monocle 3, split by time point, to demonstrate transient loss of Ciliated Progenitor cells in early infection timepoints. F,G) UMAP results for SARS-CoV-2 infected cells (blue) were (F) split by time point and (G) aligned. H) Bar graphs of the proportion of infected cells per ciliated subtype at each time point. I) Pie charts of proportion of total infected ciliated cells represented by each subtype per time point. J) Selected top gene ontology (GO) terms for differentially expressed genes (DEGs) between ciliated subtypes, plotted by significance (-log(p_val)), indicating the preferential association of viral-associated gene expression with Ciliated Progenitor cells. K) Heatmap of genes associated with the Ciliated Progenitor-associated GO term: “Viral entry into host cell” with added known and putative receptors for SARS-CoV-2, demonstrating relatively high expression of these genes in Ciliated Progenitor cells; columns represent average expression per cluster, rows are unity normalized and colored by relative expression (see Methods). L) Heatmap of expression of viral entry genes in individual ciliated cells ordered by pseudotime, and labeled by ciliated subtype and infection status, demonstrating peak viral entry gene expression and infection in ciliated progenitor cells; columns represent expression levels of individual cells, rows are unity normalized and colored by relative expression (see Methods).

With this sub-clustering, we revisited the cell population dynamics analysis. Similar to basal cells in the bulk sample, the relative number of Ciliated Progenitor cells acutely decreased at 1dpi compared to baseline (Mock), and then recovered to near-control proportions by 3dpi (Fig. 2D). This loss of Ciliated Progenitor cells was also reflected in the pseudotime analysis, revealing that the least mature cells were depleted on infection and then recovered over time (Fig. 2E). Viral load is concentrated first in Novel Infected cells at 1dpi, followed by Ciliated Progenitor cells as their numbers are replenished (Fig. 2F,G). By 3dpi, nearly 25% of Ciliated Progenitor cells are infected, as well as nearly 100% of Novel Infected Ciliated cells (Fig. 2H). By 2dpi and 3dpi, when the cell populations begin to re-stabilize, Ciliated Progenitor cells account for 60% of all infected ciliated cells, with Novel Infected Ciliated cells accounting for 28% at 3dpi (Fig. 2I).

Analysis of differentially expressed genes between Ciliated cell subtypes using GO revealed functional groupings of genes unique to each subtype (Fig. 2J). Notably, this method further suggested that Ciliated Progenitors are the main hub of viral entry and transcription among ciliated cells. GO analysis also confirmed functional roles of the other ciliated subtypes, including antigen presentation in Mature Ciliated 1 cells, cilia function in Mature Ciliated 2 cells, and interferon production in Novel Infected Ciliated cells. By pulling the genes categorized in the “viral entry into host cell” GO category, as well as known and putative genes associated with specific entry of SARS-CoV-2, such as ACE2, TMPRSS2 (Hoffmann et al., 2020), Cathepsin L (CTSL), HMGB1 (Wei et al., 2020), neuropilin-1 (NRP1) (Cantuti-Castelvetri et al., 2020), BSG (Matusiak and Schurch, 2020), and FURIN (Walls et al., 2020), we observed that most of these genes are preferentially expressed in the Ciliated Progenitor population (Fig. 2K). In addition, TMPRSS2, BSG, HMGB1, and CTSL also showed high expression in Mature Ciliated 1 cells.

Interestingly, the Novel Infected Ciliated cells, which are nearly ubiquitously infected with SARS-CoV-2, did not significantly express any genes associated with viral entry, suggesting these cells arise from already infected ciliated cells. Indeed, these cells fall just after Ciliated Progenitor cells in the pseudotime developmental trajectory. Ciliated cells ordered by pseudotime confirm that expression of viral entry genes concentrates early in the ciliated cell differentiation trajectory (Fig. 2L). These observations strongly argue that Ciliated Progenitor cells are the main point of entry for SARS-CoV-2.

### Interferon response analysis

At 3dpi, genes associated with interferon production and response are uniquely upregulated over all other time points (Fig. 3A). Comparison of genes associated with interferon-related gene ontology (GO) categories demonstrates that expression of most of these genes increases over time following infection, peaking around 3dpi (Fig. 3B). Specifically, IFIT1, IFIT2, IFIT3, and IFIT5 appear most strongly at 3dpi (Fig. 3C).

**Figure 3.**
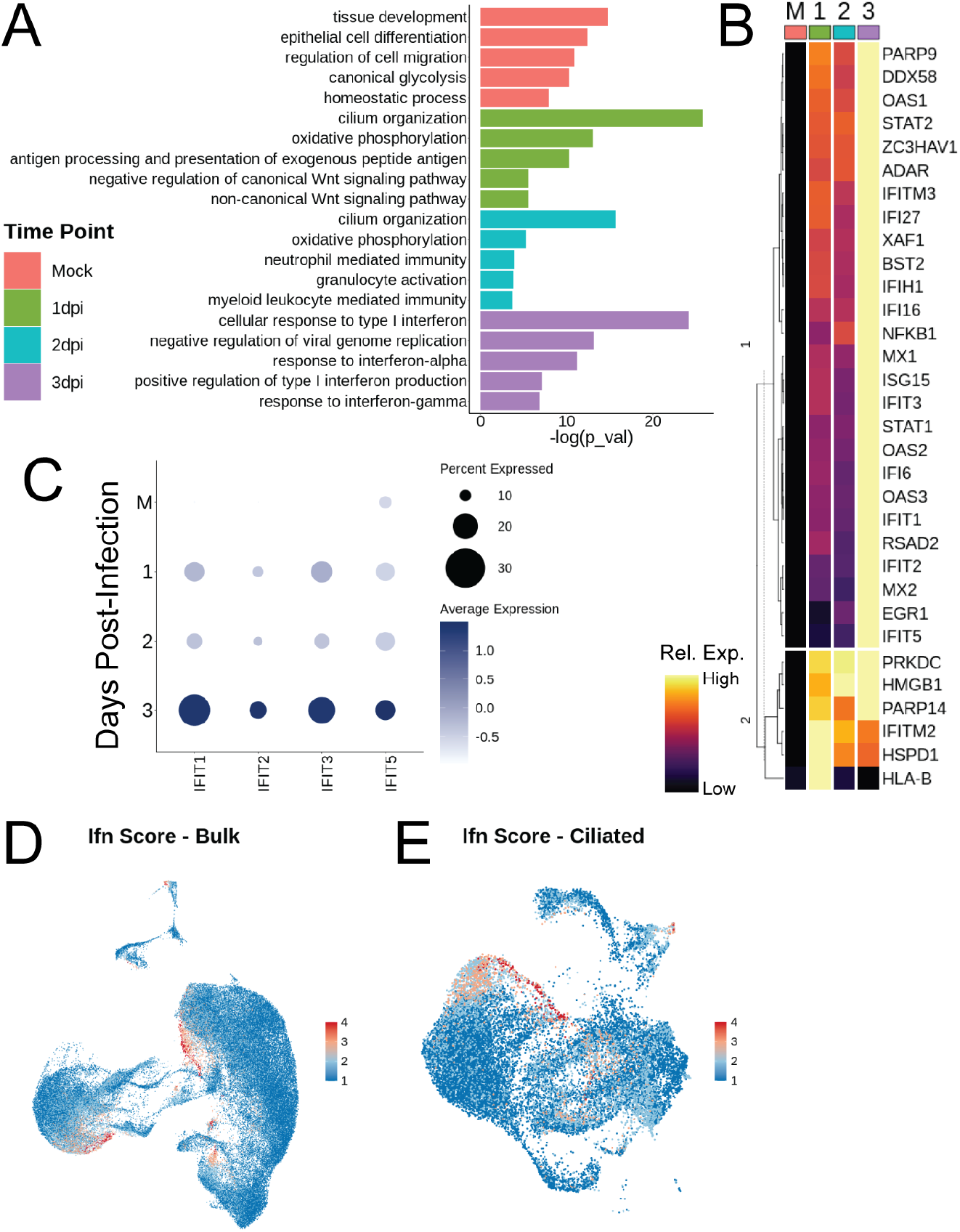
Interferon response analysis. A) Selected top GO terms for genes differentially expressed between bulk samples across time, plotted by significance. B) Heatmap of interferon gene list determined by all interferon-related GO terms in panel A, grouped by dendrogram to demonstrate expression trends over time; columns represent average expression per cluster, rows are unity normalized and colored by relative expression (see Methods). C) Dot plot of IFIT gene expression over time. D,E) Feature plots of interferon signaling score (as calculated using Seurat’s AddModuleScore with the gene list in panel (B) in (D) bulk and (E) ciliated cell populations; a high score demonstrates greater relative expression of this group of genes, suggesting interferon-related gene expression maximized at 3 days post-infection (dpi), and primarily localized to a portion of Mature Ciliated 1 cells and Club cells.

It is worth noting that these viral response signals appear as part of the innate immune response and are mounted by epithelium in the absence of immune cells (Shenoy et al., 2021). These genes mainly appear in the subsets of ciliated and club cells, which is consistent with previous findings (Chua et al., 2020), as well as some cells in PNEC, Basal, and Basal EMT populations (Fig. 3D). Interferon-related gene expression is most concentrated in an infection-specific archetype of the Mature Ciliated 1 cell cluster and in the Novel Infected Ciliated cells (Fig. 3E). Taken together, this suggests that interferon secretion is not the immediate response of epithelial cells to infection with SARS-CoV-2 virus, leaving the possibility that the epithelium may undergo another defensive response first.

### Analysis of signaling networks during infection

We then explored putative mechanisms underlying the transient enhancement of basal cell differentiation, followed by restoration of pro-renewal cues by 3dpi. Mapping the single-cell data against the FANTOM5 database of ligand-receptor interactions allowed us to explore cell-cell communication networks as a function of time following infection (Fig. 4A). We performed Wilcoxon Rank Sum tests on all genes potentially involved in intercellular communication for each major cell type shown in Fig 4B, using the Mock condition as a control for each post-infection time point(Ramilowski et al., 2015). The resultant list of all ligands and receptors displayed study-relevant and statistically significant differential expression following viral infection. Each signaling mechanism having a significant perturbation of both ligand and receptor, vs the Mock control, was then plotted over time (Fig. 4C).

**Figure 4.**
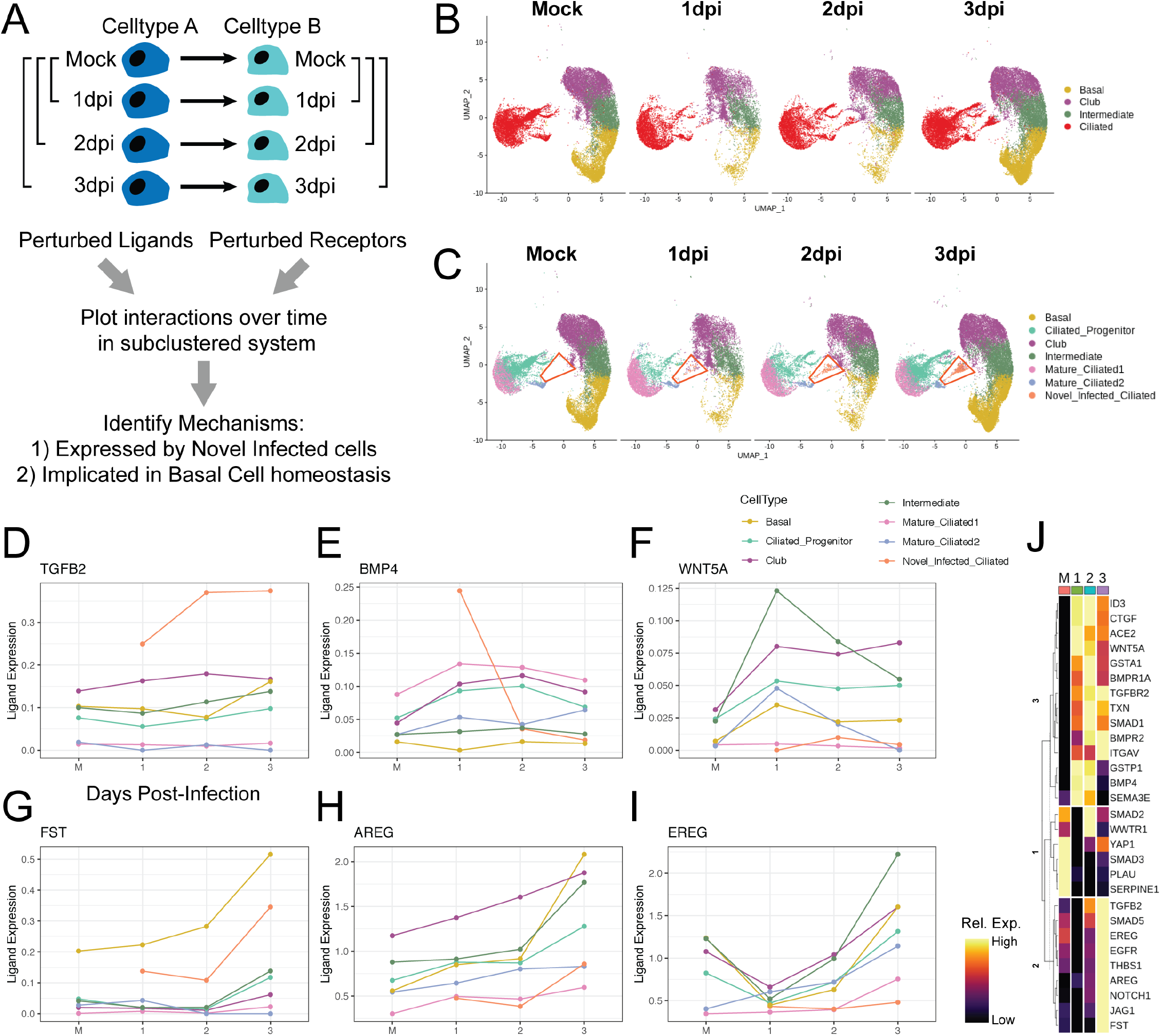
Analysis of signaling networks during infection. A) To determine the set of ligands and receptors which were perturbed by infection, paired Wilcoxon Rank Sum tests were performed, by bulk cell type, using Mock cells as the reference, for each post-infection time point on all genes in the FANTOM5 database. B) This routine was performed with the ciliated subtypes grouped as a single cell type. C) All ligand-receptor interactions found to be significantly perturbed were then plotted over time, with the ciliated population broken into subtypes; particular focus was given to the Novel Infected Ciliated cell population (red trapezoid), which does not exist in the Mock condition and only appears post-infection. D-I) Longitudinal line plots of key ligands of interest, split and colored by cell type, over time. Ligand expression was calculated using a standard log-normalization technique (see Methods). J) Heatmap of all genes of interest over time; columns represent average expression per cluster, rows are unity normalized and colored by relative expression (see Methods).

**Figure 5.**
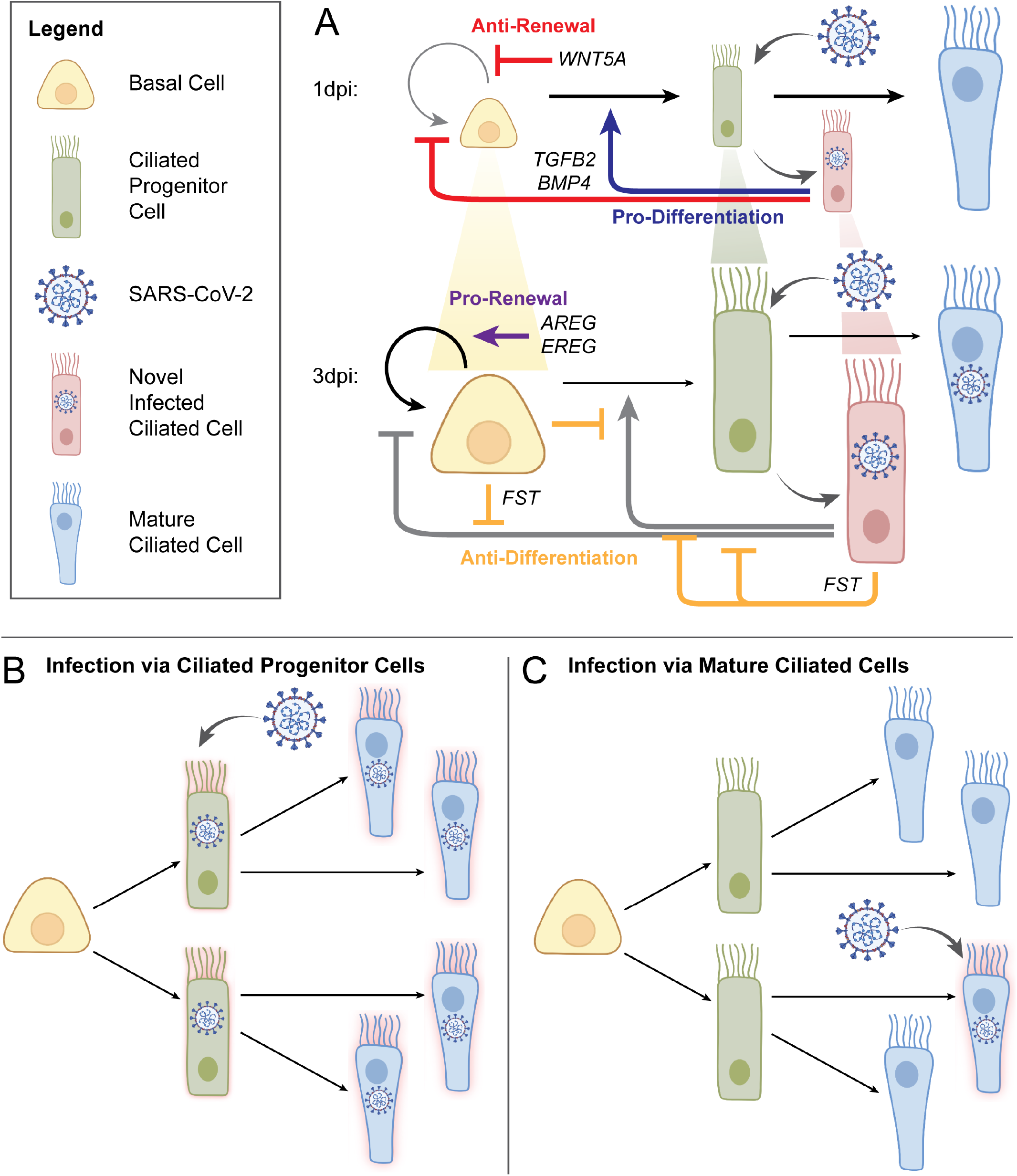
Summary schematics of cell dynamics regulated by inferred cell-cell interactions. A) Signaling mechanisms influencing population dynamics during early viral infection. At 1dpi, TGFB2 and BMP4, primarily from Novel Infected Ciliated (NIC) cells, promote differentiation and block self-renewal of Basal Cells (BC); WNT5A further blocks self-renewal in BC; this contributes to the observed depletion of progenitor cell populations, including BC and Ciliated Progenitor (CP) cells, while Mature Ciliated (MC) cells are largely maintained; increased differentiation also provides a wider window for viral entry via CP cells. By 3dpi, FST expressed by NIC and BC blocks aforementioned TGFB2 and BMP4 signaling, effectively blocking differentiation, while newly increased AREG and EREG signaling restores BC self-renewal; progenitor cell populations (indicated by cell size in the diagram) and patterns of homeostatic turnover are restored, simultaneously with upregulation of interferon-related signaling (see Fig. 3). B,C) Model of CP vulnerability to infection. Comparison of models of viral infection via Ciliated Progenitor cells (B) vs viral infection via Mature Ciliated cells (C). The virus achieves greater overall cellular infection when leveraging the rapid epithelial turnover during early infection and viral entry into CP cells, than if the virus were to enter via MC cells alone, as is frequently assumed. This summarizes our findings and demonstrates the efficiency of the viral entry strategy leveraged by the SARS-CoV-2 virus in infecting human host airway epithelium. Images adapted from BioRender.

Interestingly, we found that Novel Infected Ciliated cells, which harbor the greatest viral load, rapidly increased expression of TGFβ2 and BMP4, which can exert pro-differentiation and anti-proliferation effects on pulmonary basal cells (Fig. 4D,E) (Aschner and Downey, 2016; Bartram and Speer, 2004; Saito et al., 2018; Tadokoro et al., 2016). In addition, basal cells (and virtually all observed cell types, except Novel Infected Ciliated cells and Mature Ciliated 2 cells) expressed high and gradually increasing levels of TGFβ-receptor, TGFBR2, while maintaining high levels of BMP receptors, including BMPR1A (Supp. Fig. 1A,B). The increase in BMP4 signaling was transient, peaking at 1dpi, along with expression of its canonical target, ID3, in basal and other progenitor cells (Supp. Fig. 1C). In contrast, TGFB2 expression increased more gradually and was more persistent, along with up-regulation of TGFβ target genes CTGF, ITGAV and THBS1 (thrombospondin) (Supp. Fig. 1D-F). Interestingly, the proteins encoded by these genes have been shown to further promote TGFβ signaling, constituting putative positive feedback loops, which may explain their prolonged expression (Lipson, 2012; Mamuya and Duncan, 2012; Murphy-Ullrich and Suto, 2018).

The overall pro-differentiation signal was also likely enhanced by a transient increase in expression of a non-canonical WNT ligand, WNT5A, known to be antagonistic to canonical WNT signaling implicated in self-renewal of basal cells (Fig. 4F) (Caprioli et al., 2015; Hussain et al., 2017; Kim et al., 2019; Serra et al., 2011). However, by 3dpi, there was an increase of expression of a potent inhibitor of BMP and TGFβ signaling, follistatatin (FST), as well as a strong increase of expression of EGFR ligands AREG and EREG (Fig. 4G-I). AREG has been shown to promote basal cell self-renewal at the expense of differentiation (Zuo et al., 2017), suggesting that these signals can curtail pro-differentiation signaling. By shutting off pro-differentiation cues and encouraging self-renewal, the system can prevent total exhaustion of the basal cell pool, and support replenishment of basal and progenitor cells. Taken together, these data suggest that basal cells are initially being induced to differentiate by TGFβ and BMP signals arising from Novel Infected Ciliated cells, potentially aided by antagonism of pro-renewal WNT signaling. By 3dpi, these pro-differentiation signals are halted by inhibition of TGFβ and BMP signaling by FST, and pro-renewal of basal cells is restored by expression of AREG and EREG.

The interplay between BMP and TGFβ signaling can be complex, potentially leading to synergistic or antagonistic effects (Guo and Wang, 2009). However, our analysis indicates that both TGFβ2 and BMP4 signal through SMAD1/5 rather than SMAD2/3 signaling pathways (Supp. Fig. 1G-J). Indeed, SMAD2/3 responsive genes SERPINE1 and PLAU were downregulated in putative TGFβ target cells, in spite of the evidence of TGFβ2 activation, with SMAD3 expression also decreasing over time (Supp. Fig. 1K,L) (Zhang et al., 2011). Furthermore, we found a gradual up-regulation of genes previously implicated as direct SMAD1/5 targets, BMPR2 and JAG1, consistent with more persistent TGFβ2, rather than more transient BMP-mediated signaling kinetics (Supp. Fig. 1M,N) (Morikawa et al., 2011). In contrast, in Novel Infected Ciliated cells, PLAU expression transiently increased, suggestive of a divergent signaling program (Supp. Fig. 1L).

These observations raised the question of how the pro-differentiation, and later pro-proliferation, signals were induced. Both TGFβ2 and BMP4 can be induced in response to tissue damage from acute or chronic injury (Aschner and Downey, 2016; Bartram and Speer, 2004; Saito et al., 2018; Tadokoro et al., 2016). However, importantly, TGFβ signaling can also be induced by various viral infections (Mirzaei and Faghihloo, 2018). This could occur, for example, due to an increase of intracellular reactive oxygen species (ROS) triggered following viral entry, which has been observed in SARS-CoV-2 infection (Delgado-Roche and Mesta, 2020). We found evidence of the onset of oxidative stress in intermediate and basal cells, as indicated by an increase in expression of isoforms of glutathione S-transferase (GSTA1 and GSTP1) and thioreductin (TXN) (Supp. Fig. 1O-Q). Interestingly, TXN underwent rapid downregulation, specifically in Novel Infected Ciliated cells, suggesting a virus-mediated repression of the anti-oxidative stress mechanisms in these cells, consistent with prior results showing suppressive effects of various viral infections, including SARS-CoV-2, on Nrf2-mediated anti-oxidative stress responses (Olagnier et al., 2020). This view was corroborated by the surprising decrease of ACE2 in the Novel Infected Ciliated cells (Supp. Fig. 1R). Indeed, ACE2 has recently been implicated as a direct Nrf2 target, and thus its expression is expected to drop along with the expression of other Nrf2 targets, such as TXN (Hawkes et al., 2014; Tonelli et al., 2018; Zhao et al., 2018).

The delayed up-regulation of the basal cell self-renewal-promoting factors, AREG and EREG is of less clear origin. However, it is of interest that these genes are targets of Nrf2 and YAP/TAZ signaling (Reiss et al., 2014; Zhang et al., 2009). Even though we found that the components of the YAP/TAZ signaling network, YAP1 and WWTR1, were transiently downregulated (which could assist in basal cell differentiation (Zhao et al., 2014)), they recovered by 3dpi, which in the presence of enhanced Nrf2 signaling across different cell types could lead to elevated AREG and EREG expression (Supp. Fig. 1S,T) (Wang et al., 2020; Zuo et al., 2017).

A number of other signaling networks displayed dynamical alterations during the first few days following SARS-CoV-2 infection. In particular, we found that NOTCH1 increased gradually in basal and intermediate cells, which could further promote differentiation towards ciliated and possibly club cell lineages (Supp. Fig. 1U) (Wang et al., 2020). Trends in these and other signaling network perturbations are summarized in Fig. 4P, as well as Supp. Fig. 1.

Overall, these data suggest that a particularly high level of oxidative stress in Novel Infected Ciliated cells, not mitigated by the Nrf2-mediated response, may have led to enhanced TFGβ signaling from Novel Infected Ciliated cells to other cell targets, including basal cells. By deriving the source of this pro-differentiation signaling, we can complete the picture of the communication network that drives a brief period of rapid basal cell differentiation following viral infection.

### Putative mechanisms of viral tropism to progenitor cells due to metabolic switch

Metabolic dynamics may also be a possible mechanism of viral tropism to progenitor cells sub-populations. The respiratory pathways in the lung tissue are thought to be primarily glycolytic, which is somewhat paradoxical given the ample oxygen supply available from inspired air (Weber and Visscher, 1969). Consistent with this assumption, analysis of the Mock control displayed greater expression of genes involved in glycolysis than oxidative phosphorylation (Supp. Fig. 2A). However, subsets of cells undergoing differentiation displayed an increase in gene expression that was specific to oxidative phosphorylation, consistent with observations suggesting that oxidative phosphorylation may accompany epithelial cell differentiation (Zheng et al., 2016). Surprisingly, upon infection with SARS-CoV-2, most cells underwent a rapid switch from glycolysis to oxidative phosphorylation. This was revealed by a sustained down-regulation of glycolytic genes at 1dpi, and a corresponding up-regulation of genes associated with oxidative phosphorylation (Supp. Fig. 2A). This finding was corroborated by certain key glycolytic genes being lost on infection, along with oxidative phosphorylation genes being up-regulated (Supp. Fig. 2B,C).

Glycolysis-associated gene expression was most reduced in ciliated and basal cell clusters (Supp. Fig. 2D), whereas oxidative phosphorylation gene expression was most up-regulated in those populations (Supp. Fig. 2E). Furthermore, the cell sub-populations showing higher and increasing viral load were also the sub-populations with a particularly high oxidative phosphorylation gene expression signature (Fig. 1H). This relationship may not be coincidental, since differentiation, even under homeostatic conditions, is frequently associated with increasing oxidative phosphorylation and oxidative stress (Drehmer et al., 2016; Shyh-Chang and Daley, 2015; Tatapudy et al., 2017; Zheng et al., 2016). Oxidative stress, in turn, may promote expression of ACE2, thereby biasing viral tropism towards progenitor cells. This putative mechanism remains to be further validated experimentally, but it may help explain the apparent predilection of the virus for progenitor cells in SARS-CoV-2 infection.

## Discussion

The results presented here suggest that SARS-CoV-2 infection can trigger a rapid and transient increase in basal and progenitor cell differentiation, enabled by expression of pro-differentiation signaling cues by virally infected cells (Fig. 4). The cues are secreted by a newly described subset of infected ciliated cells (Novel Infected Ciliated cells, or NICs), that cluster separately from other mature and progenitor ciliated cells and display a virtually 100% infection rate. The pro-differentiation cues, particularly TGFβ, may be produced as a result of the increased oxidative stress precipitated by viral infection.

The process of differentiation transiently depletes basal cells, a key airway epithelial stem cell reservoir, while supporting the proportion of ciliated cells. As cellular proportions stabilize by 3dpi, up-regulation of new anti-differentiation, pro-renewal cues, particularly those of the EGF receptor family, restores the renewal of basal and progenitor cells. Although this process can serve as a typical strategy to repair the bronchial tissue damage inflicted by the initial viral infection, it may also be compromised by the apparent tropism of the virus toward active progenitor cells rather than to stable mature cells. As a result, the tissue repair process itself may open the door wider to infection by exposing increased progenitors for viral entry, and subsequent progressive infection of mature epithelial cells. A similar viral entry and infection strategy by targeting progenitor cells has been previously observed in placental infection by Zika virus and human cytomegalovirus (Tabata et al., 2016; Tabata et al., 2015).

Another interesting finding is the rapid down-regulation of ACE2 receptor in NICs. This down-regulation was correlated with suppression of other oxidative stress response genes, likely as part of the viral effect on cells. This effect may be coupled with the potential reduction in ACE2 protein content in SARS-CoV-2 infected cells (Blume et al., 2021; Glowacka et al., 2010; Verdecchia et al., 2020). A decrease of ACE2 expression may help limit the multiplicity of single cell infection by the virus, thereby increasing viral yield by the cell due to an increased cell survival by avoidance of high multiplicity of infection. This effect is observed and exploited in viral vaccine production (Aggarwal et al., 2011), and is supported by mathematical modeling (Rudiger et al., 2019). An increased viral yield, on a per-cell basis, may help spread the infection within the epithelial layers, increasing the overall pathogenicity of the virus.

These results add a new dimension to the discussion of age and gender dependence of COVID-19 progression. Indeed, children and young people may experience reduced SARS-CoV-2 infection rates due to naturally higher epithelial regenerative capacities. Homeostatic airway epithelial turnover and progenitor cell function decreases with age (Watson et al., 2020). In young people, rapid epithelial turnover in response to sensing inhaled SARS-CoV-2 particles, as observed in our study, may be adequate to successfully block viral entry or shed and replace infected cells before systemic transmission, except in cases of exposure to sufficiently high viral loads. This protective mechanism would be greatly reduced in older populations.

Furthermore, it has been suggested that reduction of ACE2 expression with age (less pronounced in females) in animal models (Xie et al., 2006), and possibly in humans may be, somewhat paradoxically, related to a lower incidence but higher severity of the disease in older patients, possibly due to an amplified inflammatory reaction (AlGhatrif et al., 2020). Our data suggest that these results may need to be viewed in a boarder context of a decrease of Nrf2 expression with aging (Zhou et al., 2018), particularly in the context of the respiratory pathways and the related increase in the oxidative stress. Enhanced oxidative stress in older individuals may lead to an enhanced expression of TGFβ and, possibly other pro-differentiation ligands (Lehmann et al., 2016; Tominaga and Suzuki, 2019), prompting differentiation of precursor cells and contributing to depletion of the progenitor cell pools. Thus, the reparative capacity of the aged bronchial and lung tissues can be compromised in the elderly. As suggested above, lower ACE2 expression in aged individuals, or in patients where ACE2 expression may be altered as a side effect of pre-existing conditions, may limit the per-cell MOI and increase the yield of viral replication and the infection spread to adjacent cells. Taken together, these effects of aging, coupled with potential age-related alterations in the immune response, may provide a wider framework in which to understand and address the age- and gender-related differences in COVID-19 severity and morbidity.

The COVID-19 pandemic is not yet beaten, and the infection dynamics of the virus must be elucidated in order to treat remaining cases and fully vanquish this disease. By identifying systemic vulnerabilities and modeling offensive viral strategies, this work may open the door to potential points of intervention for the prevention and/or treatment of SARS-CoV-2 infection.

## Acknowledgements

This work was supported by grants from the National Institutes of Health (U54 CA209992 (NCI) to A.L.; F30HL143906 and T32GM136651 to M.S.B.R.), as well as private research gifts and an unrestricted research gift from Humacyte Inc. We would like to acknowledge Drs. Craig Wilen, David van Dijk, and Akiko Iwasaki for helpful discussions and input in the interpretation of our results.

## Author Contributions

Conceptualization, A.M.G., M.S.B.R., M.P.K., and A.L.; Methodology, A.M.G., M.S.B.R., and M.P.K.; Software, A.M.G., M.S.B.R., and M.P.K.; Validation, M.S.B.R; Formal Analysis, A.M.G., M.S.B.R., M.P.K., and A.L.; Investigation, A.M.G., M.S.B.R., M.P.K.; Resources, L.E.N. and A.L.; Data Curation, A.M.G.; Writing – Original Draft, A.M.G. and A.L.; Writing – Review & Editing, A.M.G., M.S.B.R., L.E.N., and A.L.; Visualization, A.M.G., M.S.B.R., and M.P.K.; Supervision, A.L.; Project Administration, A.L.; Funding Acquisition, L.E.N. and A.L.

## Figure Legends

**Supplementary Figure 1.**
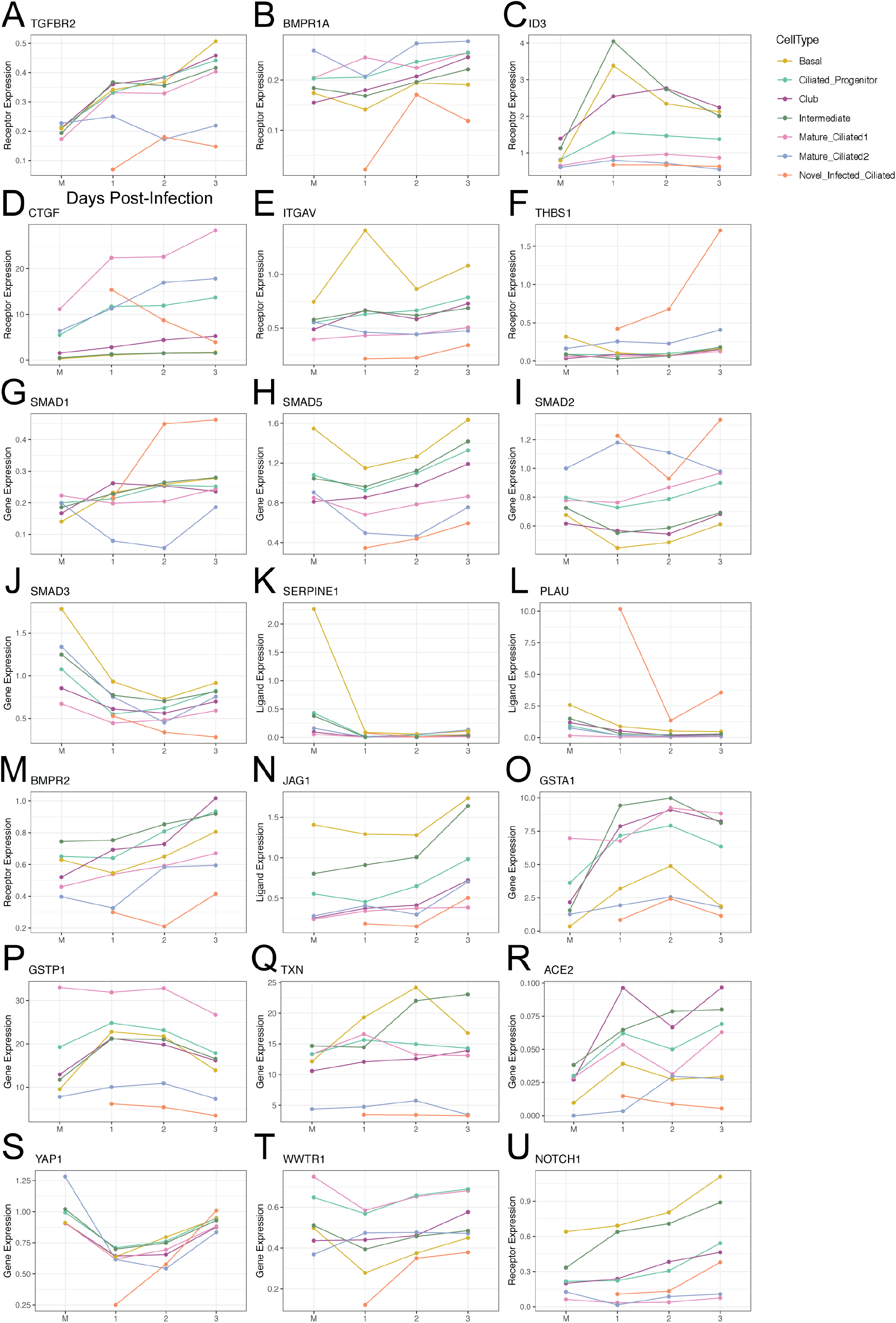
Extended analysis of ligand-receptor signaling perturbations in infected samples. A-U) Longitudinal line plots of key signaling genes of interest, split and colored by cell type, over time. All gene expression was calculated using a standard log-normalization technique (see Methods).

**Supplementary Figure 2.**
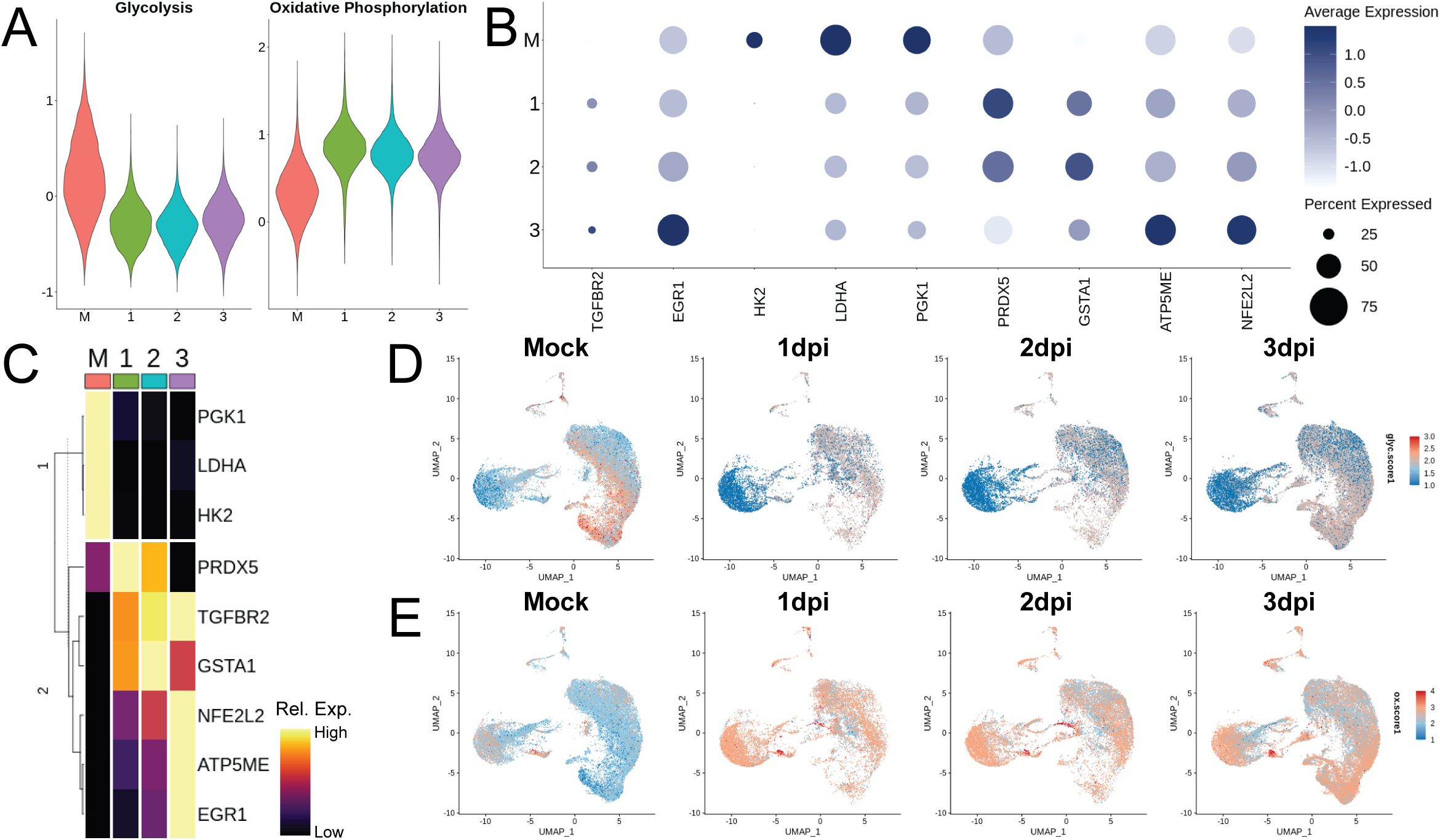
Putative mechanisms of viral tropism to progenitor cells due to metabolic switch. A) Violin plots of glycolysis and oxidative phosphorylation scores (as calculated using Seurat’s AddModuleScore with gene lists derived from associated GO terms in Supp. Fig. 1A) in bulk populations over time; a high score demonstrates greater relative expression of this group of genes. B) Dot plot of glycolysis and oxidative phosphorylation genes over time. C) Heatmap glycolysis and oxidative phosphorylation genes, grouped by dendrogram to demonstrate expression trends over time; columns represent average expression per cluster, rows are unity normalized and colored by relative expression (see Methods). Feature plots of D) glycolysis score and E) oxidative phosphorylation score in the bulk population split by time point. Overall data demonstrate a shift from primarily glycolytic to primarily oxidative metabolism with infection of cultures.

## STAR Methods

### RESOURCE AVAILABILITY

#### Lead contact

Further information and requests for resources should be directed to and will be fulfilled by the lead contact, Andre Levchenko.

#### Materials availability

This study did not generate new unique reagents.

#### Data and code availability

This paper analyzes existing, publicly available data. These accession numbers for the datasets are listed in the key resources table. All original code has been deposited at [repository] and is publicly available as of the date of publication. DOIs are listed in the key resources table. Any additional information required to reanalyze the data reported in this paper is available from the lead contact upon request.

### METHOD DETAILS

#### Generation and pre-processing of scRNAseq data

scRNAseq data of ALI cultures were obtained from Ravindra et al. (Ravindra et al., 2021). Briefly, human bronchial epithelial cells (HBECs, Lonza) were cultured at an air-liquid interface (ALI) for 28 days to achieve full mucociliary differentiation. Cultures were then challenged at the apical surface with 10^4^ plaque forming units (PFU) of SARS-CoV-2. An uninfected mock control and samples at 1dpi, 2dpi, and 3dpi were harvested with TrypLE Express Enzyme (ThermoFisher) and prepared with the Chromium Next GEM Single Cell 3’ Gel beads v3.1 kit (10X Genomics), at a target loading of 10,000 cells per sample. Libraries were generated using the Chromium Single Cell 3’ Library Kit v3.1 (10X Genomics) and sequenced on the NovaSeq 6000 using HiSeq 100 base pair reads and dual indexing, to an average depth of 31,383 reads per cell. Data were aligned and pre-processed using the 10x Genomics Cell Ranger pipeline and a combined human and SARS-CoV-2 genome. On average across samples, there were 10,000 to 15,000 counts per cell and 2,400 to 3,600 unique genes per sample. For more information, see original publication(Ravindra et al., 2021).

#### Alignment, clustering, and cell type identification of scRNAseq data

Count matrices and metadata were input to Seurat v3.0 (Satija et al., 2015). Integration was performed across infection time points using the standard Seurat v3 integration pipeline to enable consistent cell type identification(Butler et al., 2018; Stuart et al., 2019). The aligned object was further normalized, scaled, and clustered in UMAP space by PCA, regressing on percent mitochondrial reads and number of counts on the scale step. Resultant clusters were named as cell types based on unique canonical gene expression. Small clusters of doublets or partial cells were removed based on multiple cell marker expression and uniquely high read counts, or uniquely low read counts, respectively. Following cleaning and initial processing, all subsequent analyses were performed on a total of 75,245 cells across four conditions.

To further parcellate ciliated subtypes, all ciliated cells (21,510 cells) were subset and re-clustered following the same standard Seurat pipeline as above. Putative subtypes were identified and named based on functionality associated with differentially expressed genes (FindAllMarkers).

#### Downstream scRNAseq analyses

Most plotting was performed using Seurat and ggplot2 (Satija et al., 2015). All heatmaps were generated using ComplexHeatmap (Gu et al., 2016) and unity normalized by row to represent greatest and least expressive column condition, while preserving intermediate variation. Cell type and ciliated subtype population proportions were estimated by calculating the percent of each time point sample comprised by each cell population. While this method suffers potential quantitative error due to dissociation bias and capture efficiency, this error was reduced by simultaneous dissociation and GEM generation of samples. Cells were determined to be infected with SARS-CoV-2 by a previously described method(Ravindra et al., 2021). Briefly, single cells were considered infected if they contained ≥ 10 exonic SARS-CoV-2 viral transcripts, to control for viral cell surface attachment, ambient virus in the suspension, or read misalignment. Infected cell bar plots were generated by calculating the percent of each cell population at each time point that was determined to be virally infected, using the above metric for infection. Alternatively, infected cell circle plots were generated by calculating the cell population percent breakdown of all infected cells per infected time point.

Pseudotime analysis was performed using Monocle 3 (Cao et al., 2019), where cells are aligned on a dimensionless time-scale based on diffusion of gene expression. The standard Monocle pipeline was performed on the data slot of the bulk sample and ciliated subset, calling alignment arguments where applicable to control for batch effects between time point samples (Haghverdi et al., 2018). Calculated pseudotime scores were then transferred to metadata of the respective Seurat objects in order to plot on existing UMAP embeddings.

Gene ontological analysis was performed using the GO Enrichment Analysis function on the Gene Ontology web platform (geneontology.org) (Ashburner et al., 2000; Carbon et al., 2021; Mi et al., 2019). First differential gene analysis was performed (FindAllMarkers), with thresholds on min.pct = 0.5 and logfc.threshold = 0.5, among conditions of interest. These conditions included putative ciliated subtypes to differentiate functionality between clusters, and the bulk population across time points to identify global trends over the course of infection. Resultant list of DEGs was uploaded to generate lists of GO terms associated with biological processes of each condition. Top GO terms were curated to remove similar terms with comparable gene lists and irrelevant terms, and plotted by – log_10_(raw P-value). Genes associated with particular GO terms were used for various plotting and scoring. Cell scoring of expression of gene lists was performed for genes associated with interferon-related GO terms, “canonical glycolysis (GO:0061621)”, and “oxidative phosphorylation (GO:0006119)” (AddModuleScore).

In all instances where gene expression is quantified from scRNAseq data (i.e. line plots in Fig. 4 and Supp. Fig. 1), we utilized a log-normalization technique that is standard in the Seurat analysis pipeline(Satija et al., 2015). In this case, expression level is determined from the number of transcripts of a given gene measured within a single cell, relative to the total number of transcripts sequenced for that cell. These transcripts are referred to as “counts” and this relation is as follows:

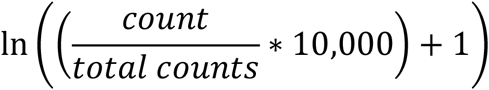

Cell-cell signaling vectors were mapped using *Connectome* v1.0.0 (Raredon et al., 2021) with a minimum ligand and receptor expression of 2.5% on sending and receiving cell respectively. Differential cell-cell signaling mechanisms were identified by performing a Wilcoxon Rank Sum test comparing populations in Mock and infected samples, for each time point, across all ligands and all receptors present in the FANTOM5 database with an adjusted p-value cutoff of 0.05. This routine was performed with the ciliated subtypes grouped as a single cell type in each sample, so as to preserve node architecture and make 1:1 connectomic comparisons possible. This provided a list of ligand-receptor mechanisms showing significant differential expression in at least one cell type-cell type vector in at least one time point comparison. This mechanism list was then used to filter the original complete connectomic edgelist to an edgelist of interest, which were considered significant. The R package ggplot2 (Wickham, 2016) was then used to create plots of longitudinal cell-cell signaling trends over time for both sending and receiving populations.

### QUANTIFICATION AND STATISTICAL ANALYSIS

#### Gene Ontology

To generate gene lists, differential gene analysis was performed (FindAllMarkers), with thresholds of min.pct = 0.5 and logfc.threshold = 0.5, among conditions of interest, to determine significantly differential genes. Top GO terms were plotted by – log_10_(raw P-value), as calculated in the GO Enrichment Analysis software, to compare significance values among terms.

#### Connectome

Significantly perturbed cell-cell signaling mechanisms were identified by performing a Wilcoxon Rank Sum test comparing populations in Mock and infected samples, for each time point, across all ligands and all receptors present in the FANTOM5 database with a minimum ligand and receptor expression cutoff of 2.5% per cluster and an adjusted p-value cutoff of 0.05.

### KEY RESOURCES TABLE

| REAGENT or RESOURCE | SOURCE | IDENTIFIER |
| --- | --- | --- |
| Deposited data |  |  |
| scRNAseq data | (Ravindra et al., 2021) | GEO: GSE166766 |
| Original code | This paper |  |
| Software and algorithms |  |  |
| Seurat (v3.0) | (Butler et al., 2018; Satija et al., 2015; Stuart et al., 2019) | https://satijalab.org/seurat/ |
| ComplexHeatmap (vX.X.X) | (Gu et al., 2016) | http://bioconductor.org/packages/release/bioc/html/ComplexHeatmap.html |
| Connectome (v1.0.0) | (Raredon et al., 2021) | https://msraredon.github.io/Connectome/ |
| The Gene Ontology Resource + GO Enrichment Analysis | (Ashburner et al., 2000; Carbon et al., 2021; Mi et al., 2019) | http://geneontology.org/ |
| Monocle (v3) | (Cao et al., 2019) | https://cole-trapnell-lab.github.io/monocle3/ |
| ggplot2 (v3.3.5) | (Wickham, 2016) | https://ggplot2.tidyverse.org/ |

